# Progressive decoding of DNA-stored JPEG data with on-the-fly error correction

**DOI:** 10.1101/2025.10.26.684640

**Authors:** Ibrahim Nawaz, Parv Agarwal, Thomas Heinis

## Abstract

DNA storage is a developing field that uses DNA to archive digital data owing to its superior information density and stability. Although DNA storage has been performed on a significant scale, challenges arise from the synthesis and sequencing of data-encoded oligonucleotides. Synthesis of DNA introduces significant noise into the process. Consequently, high-read-quality sequencers are often required, making the process expensive and lack scalability.

Error correction codes are used within the DNA storage pipeline to provide resilience to noise at a cost of additional redundancy and decoding complexity. Given such constraints and challenges, the main objective we seek to deliver is a time- and storage-efficient image coding strategy. We introduce a novel DNA-based progressive JPEG decoder with on-the-fly error-correcting & rendering capability. This system can progressively decode an image while also correcting for errors as they occur. It modifies standard JPEG encoding methods to store data in localized chunks and uses adapted Raptor error-correction codes to improve the speed and quality of partial decoding. We optimize and evaluated the method under varying levels of simulated errors, as well as show how different parts of the pipeline improve real-time decoding capability. We also test the pipeline in a real-world wet-lab experiment. We present the first progressive image decoding schema aimed at realizing real-time rendering of DNA-stored images.

## I. Introduction

Data production across the globe is growing exponentially. The International Data Corporation (IDC) predicted that globally data would increase from 45 to 175 zettabytes between 2019 and 2025 [38]. Moreover, it is expected that the storage supply will not be able to meet such a demand. By 2030, storage shortages will account for two-thirds of demand [39]. Therefore, there is a growing need to find more sustainable and efficient data storage mediums. DNA has been proposed as a suitable candidate, especially for archival purposes. DNA is extremely information dense: a gram can theoretically encode 455 exabytes of data [16]. Solid-state hard drives and magnetic tape are far less optimal. DNA is also very stable, lasting hundreds of years at room temperature or potentially millions of years in permafrost [46]. Data stored on hard drives and tape can only last for decades. DNA is also easily replicable with very low energy operational requirements. Proof-of-concept works have been conducted that encode digital data in DNA for use in storage [9], [17]. The field has since progressed significantly, from encoding a 23-byte message [8] to 200.2MB of unstructured data [35].

Storing data in DNA involves encoding information into sequences of oligonucleotides (oligos), followed by synthesis and physical storage. Retrieving the information involves sequencing the oligos and decoding them back into digital data. The main challenges to DNA storage arise from synthesis and sequencing of the oligos: alongside increased cost, they both introduce significant noise into the workflow. Noise in DNA storage usually occurs as insertion-deletion-substitution (IDS) errors (where a base is incorrectly added, removed, or replaced, respectively) or erasures (where entire oligos are dropped) [24]. Synthesis, polymerase chain reaction (PCR) amplification, and storage conditions can also cause oligos to decay at different rates [20]. Previous works [7], [18] heuristically defined some key biochemical constraints to minimize such errors when encoding and decoding: 1) Strands below 200 nucleotide (nt) length; 2) No homopolymers (repeats of a base, e.g. “TTT”); 3) GC content between 40-60%; 4) No repeated patterns in an oligo (repeated sequence of bases, e.g. “ACTACTACT”). Synthesized oligos are also stored in an unstructured fashion, which necessitates the need for extra indexing information within the oligo [18] and increases erasures when sub-sampling from the oligo pool for sequencing & decoding [21].

Prior works have shown that errors can be corrected to recover data if appropriate encoding schemes are used. Error-correcting codes introduce redundant information when encoding, allowing for erroneous data to be recovered [45] at a cost of increased storage size. Simple techniques such as parity checking [18] and overlapping segments [7] resulted in unrecoverable gaps due to error, even after using expensive high-fidelity synthesis and high-accuracy Illumina sequencing. Other studies deployed Reed-Solomon (RS) codes [5], [19], [34] which achieved full data recovery upon decoding, though were still far from realizing channel capacity. They also suffered when scaling to larger files due to variations in erasures of different oligos. Such challenges can be solved with more complex encoding schema & higher sequencing coverage. Fountain codes, like Luby Transform (LT) codes [28], are near-optimal rateless erasure codes and are able to approach the DNA storage channel capacity while providing greater robustness against erasures [14]. Modern fountain codes like Raptor [44] & Online codes [31] recently showed significantly better performance when compared with LT coding [42]. While previous literature greatly improved encoding efficiencies, they lacked scalability due to use of expensive high performance synthesis & sequencing techniques.

More scalable methods have been explored: for example using photolithographic synthesis [3] and Oxford Nanopore Technologies (Nanopore) sequencing [29]. Such techniques are more error-prone, resulting in more complex coding strategies and less efficient codes. For example, performing clustering [3] or iterative alignment [50] prior to decoding can leverage high coverage to reduce errors. However, these methods are also slow.

Our work revolves around image encoding. Image-specific encoding methods for DNA storage have also been explored. Previous works have encoded quantized wavelet coefficients of an image into DNA with a controllable compression & oligo length value [11]; implemented a JPEG-style encoding scheme [12], leveraging ternary Huffman tables [18] to replace the Huffman encoding used in JPEG. JPEG-DNA [23], currently in development, implements Raptor encoding [42] of a JPEG image. Due to inherent image compression, such methods require all information to be correctly decoded before rendering is possible. Progressive JPEGs [47] can render an image at reduced resolution before retrieving all image coefficients, and have not been explored in DNA storage before. Although developed primarily for rendering images in poor network conditions, this feature allows us to address the slow nature of decoding. Progressive JPEGs split up an image into separate scans containing different levels of detail: spectral selection [2] initially encodes lower frequency coefficients from discrete cosine transform (DCT) and iteratively encodes increasing frequency coefficients in subsequent scans. JPEGs can also use restart intervals (RI) to provide error resilience to the internal state of the JPEG decoder, and have not been explored in DNA storage before. In file system fragmentation, RIs have been used to reduce file fragmentation effects when recovering corrupted JPEG images [43].

Taken together, these results inspired us to seek a fast & scalable coding strategy with on-the-fly rendering and partial decoding. To reduce the time overhead during sequencing & decoding, we wish to utilize the progressive capabilities of JPEG to iteratively render an image as oligos are being sequenced. Further recovery will add further detail to our rendered image. Previous works have applied JPEG-specific encoding methods to DNA storage [25]. However, this is the first work to implement progressive JPEG rendering with error correction into a DNA storage workflow.

## II. Methods

### A. Custom JPEG encoding

A 680 × 453 uncompressed RGB bitmap image was encoded into a progressive JPEG using the libjpeg-turbo library [26]. We follow the typical progressive JPEG encoding schema which includes an initial scan for all DC coefficients followed by three successive AC coefficient scans for the luminance (Y) and chrominance (Cb, Cr) components [22]. The quality parameter was fixed at 75 (default) and chroma subsampling was disabled. During JPEG encoding, coefficients are compressed into variable-length Huffman codes and undergo bit stuffing when encoded into the Entropy-Coded Stream (ECS). In typical JPEGs, Restart Interval (RI) markers can be inserted into the ECS after a user-defined number of image blocks. This user-defined number is stored in the header of a JPEG file, while the markers themselves are inserted directly into the ECS, as shown in Fig 1.

**Fig. 1:**
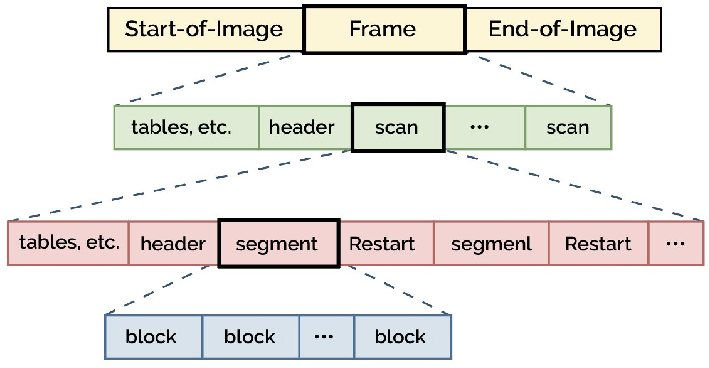
JPEG bitstream [32]. Multiple scans of coefficients form the progressive JPEG. Each scan contains segments (ECS) with restart markers (RIs) inserted at regular intervals to inform the decoder of its position in the image.

These markers have a 2-byte size and range from `0xFFD0` to `0xFFD7`. They are inserted in a sequential order from `0xD0` to `0xD7`, before restarting back to `0xD0`. Fragmentation during DNA decoding creates gaps in the ECS, necessitating the need to store global location information about RIs. Thus, RI markers were translated into ‘global’ RI markers to act simultaneously as restart intervals & as image block indices, by storing a global RI count in base-8. For example, the 10th RI marker in a scan would be converted from [`0xFF`, `0xD1`] (2 bytes) to [`0xFF`, `0xD1`, `0xFF`, `0xD1`] (4 bytes). Eq (1) shows how the RI block step size, RI_step_, was calculated dynamically, based on the required Raptor chunk size:

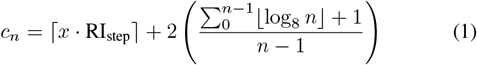

Where 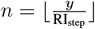, *x* = average block size, *y* = total number of blocks.

RI_step_ was iterated until *c*n < *s* · *c* (scaling factor · required chunk size) and was returned. *s* was optimized heuristically. RI_step_ itself was optimized for only the first DC scan, to minimize overhead & maximize progressive decoding when rendering. Therefore, a compressed DNA-adapted JPEG image (.jpg) was generated from the original uncompressed bitmap image (.bmp), with 4 scans and an optimized restart marker step size, RI_step_.

A JPEG decoder was implemented with the functionality to translate the global base-8 RI markers into block index information. The header data, accounting for ~5% of information, was stored off-DNA to simplify the workflow. Partial decoding involves recovering random chunks of the bitstream from random oligos. Therefore, the JPEG decoder was also adapted to validate a scan before attempting to decode it. A scan was determined as valid if it contained any span of bits starting from an RI marker (`0xFFD0`–`0xFFD7`) while including any coefficients within a recovered chunk. These spans can then be decoded independently from each other due to Huffman’s prefix coding.

### B. Adapted Raptor coding

Reading DNA-stored data involves selecting a subset of encoded oligos to sequence from an unstructured pool. This resembles an erasure channel, where data is irrecoverable until enough strands have been sequenced. This is where modern fountain codes shine and were therefore selected. The Raptor encoding schema used in JPEG-DNA [33], [42] was adapted to generate oligos from the compressed DNA-adapted JPEG image described above. The bitstream was split up into *k* equally sized chunks, according to a user-selected chunk size, *c*; *m* auxiliary chunks were generated using Gray & low-density parity-check precoding [1], [15]; a Mersenne Twister algorithm [30] was used to randomly select *d* chunks from the (*k* + *m*) original chunks. The algorithm uses the seed of a pseudorandom number generator, which models the Raptor (Robust Soliton) distribution, to sample the degree, *d*, of the packet. The packet was generated by XORing the *d* selected chunks together and prepending the seed ID. Fig 1 shows the structure of a candidate oligo packet, after bit-to-DNA translation at the entropy limit of 2-bits-per-nt.

Given a specified chunk size, *c* the oligo length can be calculated by Eq (2):

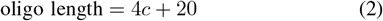

Note that alongside the Raptor-encoding depicted in Fig 2, we add a 4-nt checksum. The final step enforced previously detailed biochemical constraints. Fountain codes like Raptor are rateless, meaning they can generate an unlimited number of encoded packets. Therefore, if a candidate oligo was invalidated by the constraints, that packet was dropped, and a new one was generated. This process was repeated until *k*(1 + *θ*) valid packets were generated, where the user-defined overhead, *θ*, was set to the default value of 0.4. This value was fine-tuned based on the channel characteristics. Finally, the array of valid encoded packets was saved into a FASTA file.

**Fig. 2:**
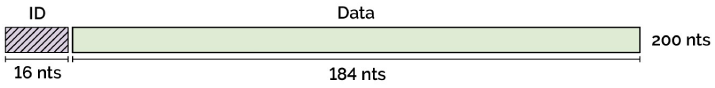
Raptor-encoded oligo packet. A 200-nt oligo generated from a 50-byte Raptor-coded packet, containing 4 bytes ID & 46 bytes data.

The Raptor decoder utilized Gaussian elimination with partial pivoting (GEPP) rather than belief propagation [40], owing to its deterministic nature, numerical stability, and predictable performance. Performance of partial decoding is expressed as the normalised area under the recovery curve, or Recovery Efficiency (RE), given by Eq (3):

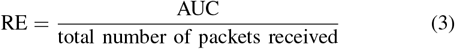

where the area under recovery curve (AUC) is calculated using the trapezium rule.

Traditional progressive JPEGs render scans in a sequential nature, where lower frequencies are rendered first. However, Raptor decoding is random – there is no control over which chunks in the ECS are retrieved first. To tackle this problem, we implement prioritized LT codes [49]. Prioritisation occurs by ensuring low-degree (1 or 2) packets have a higher likelihood to contain high-priority chunks during encoding. This allows them to be recovered faster during Raptor decoding, with negligible overhead. We assigned chunks containing the first DC scan as high-priority.

Finally, Mersenne Twister was adjusted in order to evaluate the effect of increasing degree-1 packets on partial decoding performance and overhead. A pseudorandom number was selected, *p* ∈ [0, 1]. If this value was below the degree-1 threshold (i.e. *p* < *p_thr_*), *d* was set to 1. Otherwise, the original Raptor distribution was used. We calculate the final overhead, *ε*, as shown by Eq (4):

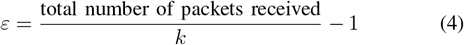

### C. Nanopore sequencing & clustering

Nanopore sequencing was simulated with Badread [48] (v0.4.1, https://github.com/rrwick/Badread) integrated with an agglomerative clustering algorithm. Similar to previous clustering-based decoders [3], [35], our clustering algorithm loosened invalidity & coverage constraints, by aggregating noisy sequence reads into clusters and using the Levenshtein distance as a similarity metric [24]. The “nanopore2023” error model was used to simulate ONT R10.4.1 (2023).Read identity was modeled by a beta distribution parameterized by mean, maximum, and standard deviation (*µ*, max, *σ*). To assess the impact of sequencing performance on overhead, we varied the mean and maximum read identity values. Sequencing depth was also varied to evaluate its effect on strand recovery during clustering. All other Badread parameters (junk reads, random reads, chimeras, adapters) were set to default values.

The clustering workflow began by grouping noisy reads from the FASTA oligo pool into strand families, where each family represented repeats of the same oligo. Reads were first filtered to retain only those within a valid length range, defined as ±5 edits from the expected strand length (accounting for adapter sequences). Adapters were then removed using k-mer search to identify and trim adapter sub-sequences.

Agglomerative clustering was applied to iteratively group strands: a read was assigned to an existing cluster if its Levenshtein distance to the cluster head was ≤ 40 edits; otherwise, it initiated a new cluster. Clustering continued until the number of clusters matched the expected number of encoded packets (determined by chunk size *c* and overhead *θ*). Consensus sequences were generated from each cluster by randomly sampling 15 reads, performing multiple sequence alignment with MUSCLE [13], and applying majority voting. A 4-nt checksum appended to each Raptor-encoded oligo served as a parity check to validate consensus candidates. This clustering approach averaged out insertion, deletion, and substitution (IDS) errors, while the checksum ensured candidate validity.

By incorporating this clustering algorithm, the storage channel could be modeled as an erasure-only channel with erasure rate *ϵ* given by Eq 5:

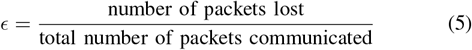

The erasure rate informs the required encoding overhead *θ* as shown in Eq 6:

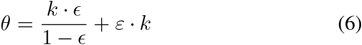

To simulate ONT’s ReadUntil functionality [27], which enables real-time sequence readout, strands were fed iteratively into the decoder. The complete pipeline is illustrated in Fig 3.

**Fig. 3:**
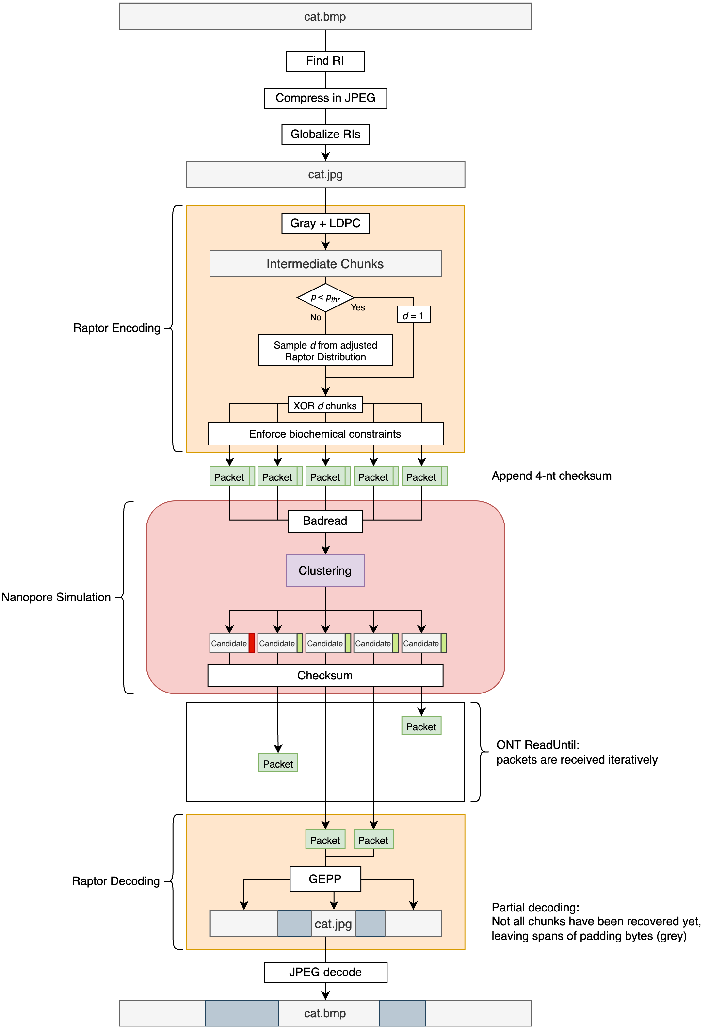
The full pipeline. Schematic of the JPEG + Raptor pipeline. Packets are fed iteratively into the decoder to simulate real-time sequencing.

### D. Experimental validation

The full pipeline was validated using synthesized DNA oligos. In traditional code analysis, the code rate 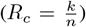 defines the efficiency. Rateless codes like fountain codes can generate an unlimited number of encoded packets. As *n* tends to infinity (unlimited number of packets), the code rate approaches 0. Practically, the number of packets generated (and therefore *n*) is fixed by a user-defined parameter, *θ*. Finding this value involves analysis of the code and channel; fountain codes therefore focus more on overhead rather than code-rates. Given an estimated model of communication, the overhead, *ε*, can be calculated. This provides a lower bound on *θ*, the number of packets to be generated. Finding this value allows us to estimate packet requirements when synthesizing. Results from this in vitro experiment are presented below in Section III-F.

### III. Results

### A. Globalising RIs

Fig 4 demonstrates the trade-off between oligo length and file size overhead when incorporating global base-8 restart interval markers during JPEG compression of cat.bmp into cat.jpg, calculated using Eq 1. The plot reveals three distinct regimes: for short oligos (between 100 and 150-nt), file size increases are substantial, reaching up to 150% for *s* = 0.33, 80% for *s* = 0.50, and 38% for *s* = 0.66. In the intermediate range (between 150 and 250-nt), all three spacing parameters show exponential decay in overhead. For longer oligos (*>* 300-nt), the file size increase approaches a plateau of approximately 10-15% for *s* = 0.33, while *s* = 0.50 and *s* = 0.66 converge to minimal overhead (around 5%). This behavior reflects the diminishing impact of fixed-size RI markers as the payload-to-marker ratio increases with oligo length. The step-like transitions in the curves correspond to discrete changes in the number of RI markers required per oligo as length increases.

**Fig. 4:**
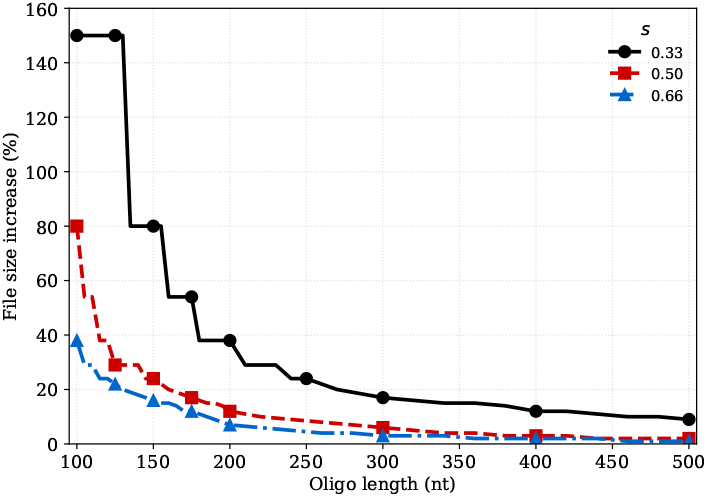
cat.jpg global RI size increase. File size increase (compared to naive JPEG) across different restart interval scaling factors.

Setting *s* = 1.0 inserted RI markers at intervals equal to the chunk size *c*, which significantly increased the likelihood of markers spanning chunk boundaries. This fragmentation of the global index resulted in incorrect pixel rendering positions. Setting *s* = 0.66 allowed for a balanced file size increase against fragmentation avoidance.

### B. Degree-1 thresholds

Changing the degree-1 threshold, *p_thr_*, affects the profile of recovery. A lower *p_thr_* increases the probability of obtaining a lower degree packet, helping to initiate the decoding process. Fig 5 compares *p_thr_* = 0.0 (i.e., the original Raptor distribution) with *p_thr_* = 0.25 & *p_thr_* = 0.5 in a noiseless scenario (*c* = 46, *k* = 619). Higher values of *p_thr_* require more received packets to achieve complete recovery but provide substantially better intermediate decoding performance throughout the process.

**Fig. 5:**
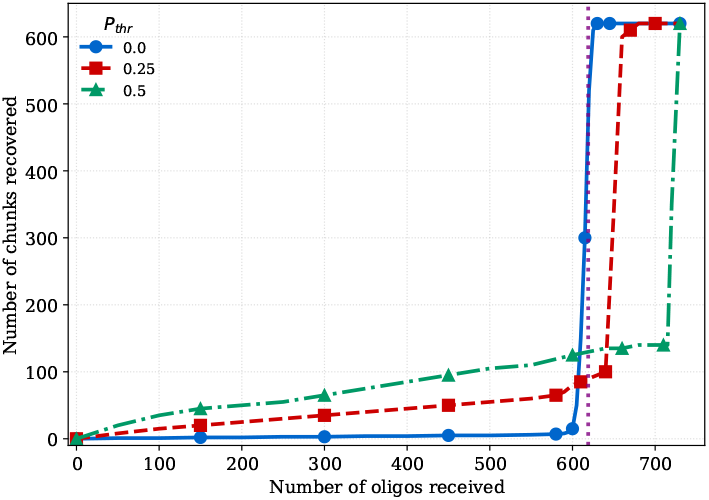
Affect of degree-1 threshold on recovery profiles. c = 46 bytes (200-nt); k = 619 (purple line).

Fig 5 also illustrates the characteristic phase transition from minimal initial recovery to rapid complete decoding. The dotted vertical line at *k* = 619 packets serves as a reference point for this transition behavior. Modifying the Raptor distribution parameters increases the decoding overhead, defined as the additional packets required beyond the theoretical minimum of *k* packets to achieve full recovery. In the noiseless case with standard Raptor parameters (*p_thr_* = 0, blue line), the overhead approaches zero [44], resulting in the most pronounced phase transition where recovery remains minimal until near *k* packets, then jumps sharply to completion.

Fig 6 shows the progressive decoding of cat.jpg with partial decoding, via GEPP. We simulate real-time sequencing by iteratively reading packets into the decoder.

**Fig. 6:**
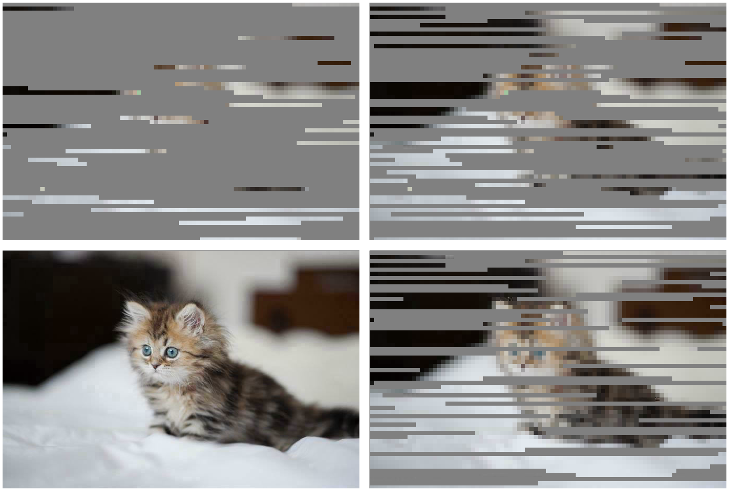
Progressive decoding while iteratively receiving more packets. Clockwise from top left: 130, 316, 660 and 693 oligos received (*c* = 46, *k* = 619).

### C. Effect of chunk size

To quantify the improvement in intermediate recovery performance, we calculate the AUC normalized by the total number of packets received to account for varying overhead. We employ Recovery Efficiency (RE) rather than simple averaging due to its superior characterization of the final recovery phase. Fig 7 demonstrates that increasing *p_thr_* enhances RE, while increasing *c* reduces RE.

**Fig. 7:**
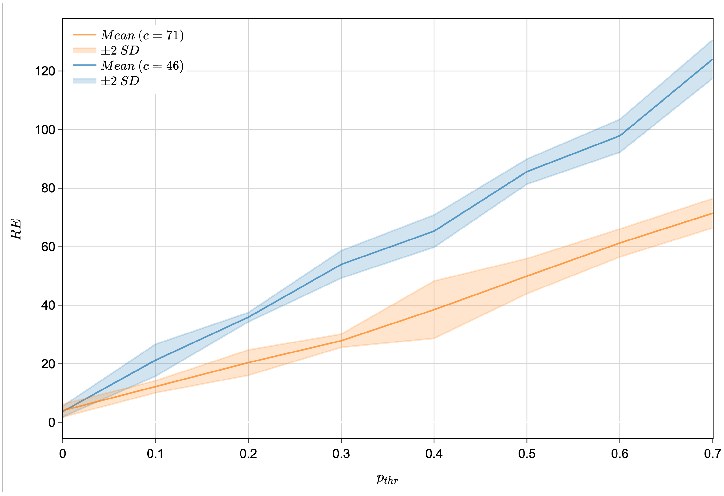
Recovery Efficiency and *p_thr_* across different chunk sizes. c (blue) = 46 bytes (200-nt); c (orange) = 71 bytes (300-nt); number of runs = 5.

Fig 8 shows the relationship between *p_thr_* and decoding overhead (*ε*) for two parameter settings. Both conditions demonstrate an exponential increase in overhead as *p_thr_* increases. The error bands (±2 SD) indicate increasing variability in overhead measurements as *p_thr_* increases, particularly beyond *p_thr_* = 0.5. Both conditions show minimal overhead (*ε* < 0.1) for *p_thr_* ≤ 0.4, indicating that moderate increases in the degree-1 threshold incur relatively small penalties in decoding efficiency.

**Fig. 8:**
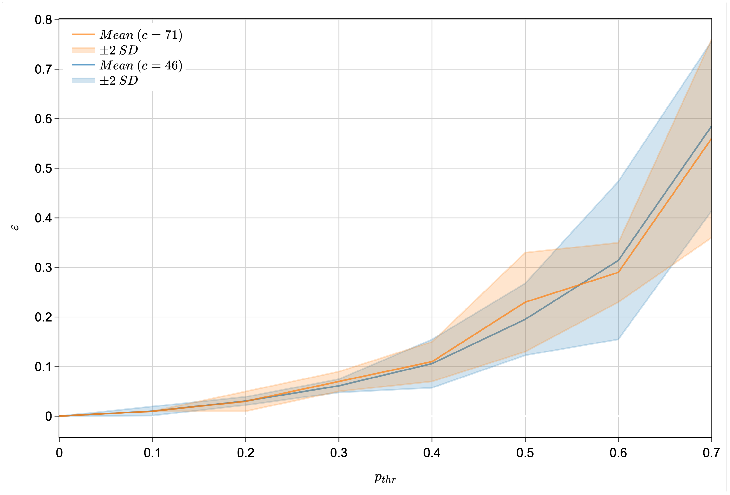
Overhead and *p_thr_* across different chunk sizes. c (blue) = 46 bytes (200-nt); c (orange) = 71 bytes (300-nt); number of runs = 5.

### D. Priority Raptor codes

Lower degree packets help initiate the decoding process in a cascading fashion. As a result, chunks stored in these packets are typically successfully decoded first. Fig 9 compares priority encoding with non-priority encoding, showing that DC scans (rather than random chunks) are recovered first when prioritized encoded. By prioritizing chunks in this fashion, we can control the information that is decoded first, which is necessary to load the image in a progressive fashion.

**Fig. 9:**
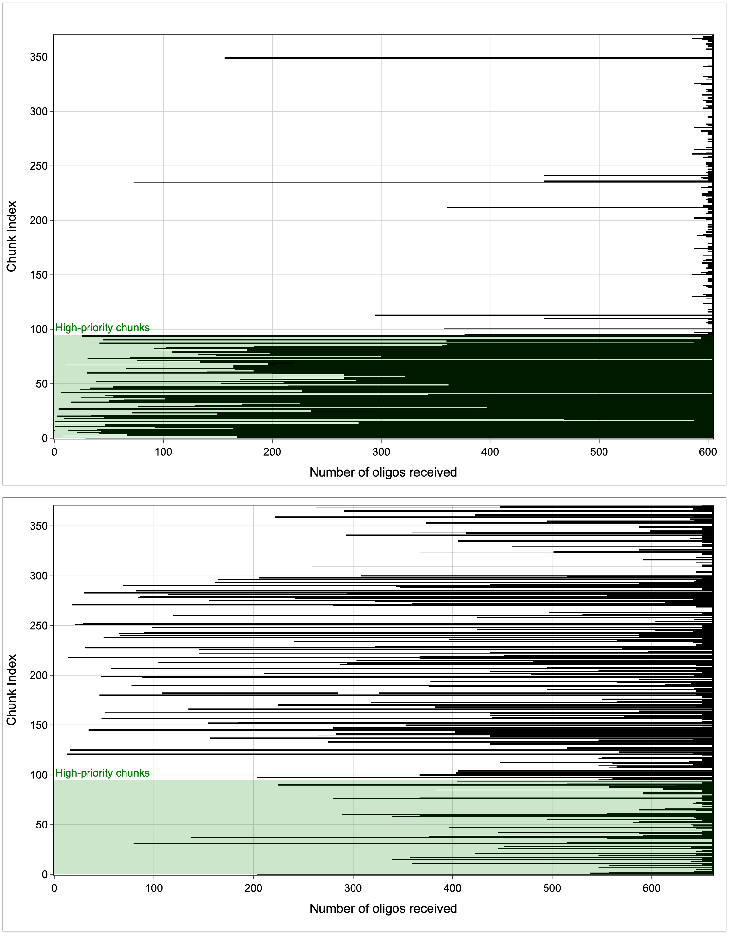
Priority vs. non-priority Raptor coding. Recovery of individual chunks with high-priority (top) and naïve Raptor encoding (bottom). Black indicates a chunk is successfully recovered. The green shaded area denotes high-priority chunks, comprised of chunks containing the DC scan (*c* = 71, *k* = 371, *p_thr_* = 0.7).

### E. Nanopore modelling

Simulating Nanopore reads allows us to both test our clustering algorithm and determine the erasure rates of the overall channel. Fig 10 shows the effect of depth on the erasure rate, with a fixed read identity Badread error model, while Fig 11 shows the effect of read identity on the erasure rate, with a fixed depth.

**Fig. 10:**
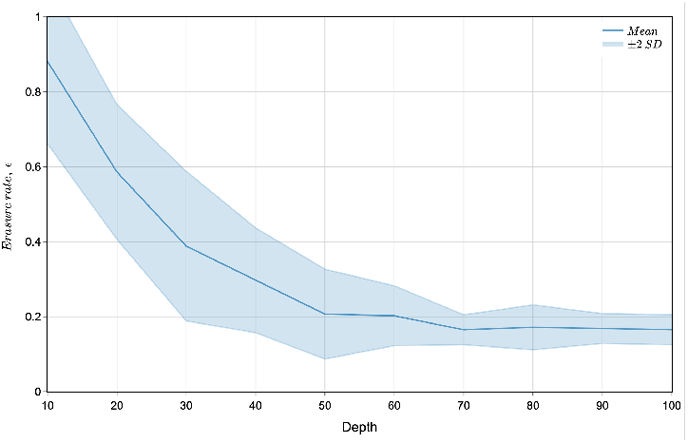
Varying depth affects the erasure rate. Read Identity = (97, 99, 2.5); *c* = 47; *p_thr_* = 0.4; number of runs = 5.

**Fig. 11:**
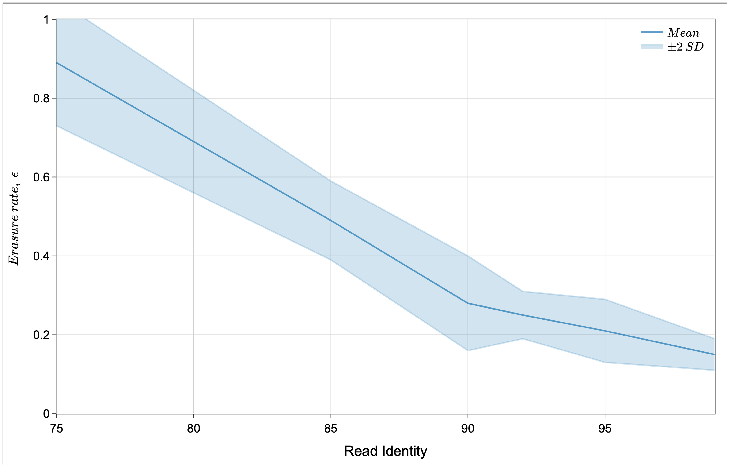
Varying read identity affects erasure rate. Depth = 40x; *c* = 47; *p_thr_* = 0.4. Read Identity max and *σ* remain fixed; number of runs = 5.

When depth *>* 60x, the erasure rates stabilize to around 0.2. Therefore, using Eq 6 we can calculate the required overhead, *θ*: around 212 strands. This provides an approximate minimum overhead when encoding at that configuration. A higher depth increases the accuracy and reliability of reads but is not scalable. Lowering depth increases the chance of error, especially insertion & deletion errors. Sequencing technologies like Nanopore, also have biasing towards certain strand patterns [10]. Badread applies depth to oligos in a non-exact fashion to simulate similar behavior. Increasing depth reduces the effect of these errors & bias. Depth = 60x gives us a good balance of both performance and cost.

### F. Wet lab experiment

Based on the Nanopore simulation results, we find *ε* to be around 0.2, and set *θ* to 0.5 for redundancy. Setting *c* = 47 and *p_thr_* = 0.4, we generate a FASTA file from cat.jpg containing 852, 208-nt length oligos (*k* = 606), using *θ* = 0.5.

These strands were synthesized by Twist Bioscience. Forward and reverse primers (20-nt each) were added to the strands prior to PCR amplification to generate a large strand pool containing multiple copies of each strand. The amplified strands were sequenced using a Nanopore sequencer (MinION MK1C), basecalled and then subsequently clustered with varying strand counts per sample. Fig 12 presents the recovery profiles for both chunks and packets as a function of the percentage of strands from the pool that were clustered. Packet recovery shows linear growth with increasing strand percentage, whereas chunk recovery demonstrates profiles consistent with the simulated results presented in Fig 5. Due to the shift in the *p_thr_*, intermediary decoding performance is improved, allowing us to recover and render the prioritised DC scan chunks first.

**Fig. 12:**
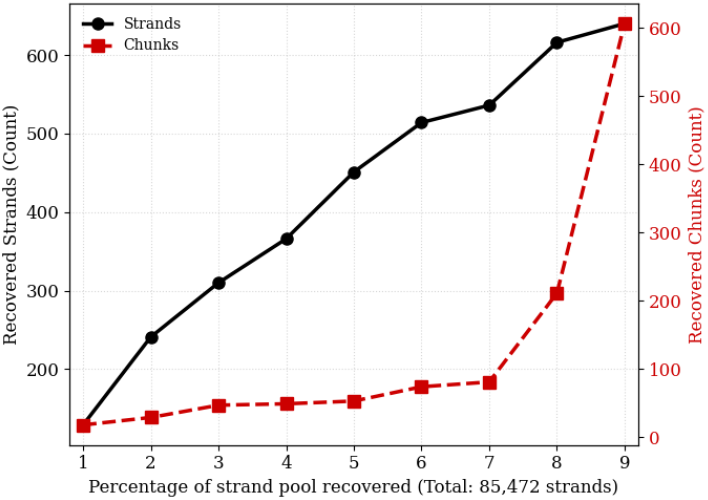
Synthesised oligo recovery profiles. Recovery profiles for packets and chunks obtained from synthesized oligos, shown as a function of the proportion of the strand pool recovered.

Due to the short nature of the checksum (4-nt), the pipeline was prone to false-positives when tested *in vitro*. 2 strands had to be heuristically removed for the decoder to finish solving. A more robust parity check scheme (CRC32) must be used in the future to avoid this.

## IV. Discussion

Our implementation adapts both JPEG and Raptor encoding. Major contributions include the integration of a fragmentation-resilient JPEG decoder via global RIs, improved intermediary decoding performance leading to enhanced recovery efficiency via a controllable degree-1 threshold and traditional progressive image decoding from low to high frequency coefficients via priority Raptor (LT) codes. While the pipeline leverages the properties of Raptor and JPEG to enable progressiveness, there are key constraints.

The observed phase transition severely limits intermediate decoding ability, and is a quality of near-optimal codes such as fountain codes. These codes achieve near-optimal (i.e., no overhead) performance by distributing information bits across packets. This has a de-localizing effect on information and packets, making them more resilient to error: each packet is less critical to the decoding process. This property has an effect of diminished partial decoding performance. A certain number of packets need to be retrieved before significant progress can be made during decoding.

Traditional Gaussian elimination (including GEPP) requires full rank systems before solutions can be calculated. Alternative methods to enable partial decoding include generalized inverse [4], Gauss–Seidel method [41], belief propagation [36], and partial Gaussian elimination [44]. We implement the latter, by iteratively eliminating as far as we can find a pivot. Once we reach a non-pivot row, we stop and attempt to solve the successfully eliminated rows. This allows us to find solutions even in an underdetermined linear system. Some chunks remain unrecoverable until the system reaches full rank, beyond which significant decoding is possible. This accounts for the sharp sudden rise of recovered chunks [6].

The rows of the system’s binary coefficient matrix correspond to a received encoded packet; columns correspond to original source chunks. Element values correspond to whether that chunk was used when generating the packet: 1s denote the presence of a chunk, and 0s denote the absence of a chunk. Thus, the degree distribution is also directly related to the sparsity of the coefficient matrix. Fig 7 shows the effect of increasing sparsity on partial decoding performance. A sparse system also reduces the effect of fill-in and increases row independence [37]: rank deficient matrices perform much better with increasing sparsity. These benefits are magnified in a binary matrix.

Degree-1 packets are constructed of one chunk, making them essentially uncoded data. These packets form the starting point of decoding, as we use these packets to reduce higher-degree packets (from *d* = 2 packets, and so on). Therefore, a higher proportion of degree-1 packets can benefit partial decoding and progressive JPEG decoding greatly. Usually, the feasibility of increasing degree-1 packets is limited by the noise in the DNA storage channel. However, our clustering algorithm is able to reduce the effects of such noise; finding a suitable *p*thr is key.

The pipeline is able to progressively render JPEGs from lower to higher resolutions by encoding low frequency scans in the prioritized chunks. Priority encoding shows great performance even with a non-zero *p_thr_*, and interestingly with minimal effect on the overhead. Once the coefficient matrix reaches full rank, recovery strongly increases. Priority encoding is what enables our pipeline to render progressive JPEGs with low frequencies first (as shown in Fig 6).

Our pipeline provides a novel image-based DNA-storage schema which is capable of rendering JPEGs progressively, with very little overhead. Excluding sequencing erasures and setting *p_thr_* = 0.4, we only introduce around 10% overhead to do so. A similar rendering effect with uncompressed images would perhaps have a lower overhead, but with a much larger oligo pool size. For example, encoding the original uncompressed cat.bmp would require around 35 times more DNA.

Clustering condenses IDS errors into erasures, at a cost of code rate. Our clustering algorithm stabilizes partial decoding and thus progressive decoding. We experimented with inner RS codes applied at an individual oligo level, but suffered from poor decoding performance. In most cases, errors exceeded the RS code’s capability. Furthermore, increasing the number of RS repair symbols increases the overhead greatly. Therefore, we mitigated IDS errors by implementing a Levenshtein distance-based clustering algorithm. Filtering with a 4-nt checksum ensured candidates were error-free, and reversed strands were reoriented. As seen in the wet lab experiment however, a stronger parity check is necessary for the decoder to function appropriately.

In summary, we propose a DNA-based progressive JPEG coding schema, with on-the-fly error-correcting & rendering capability. We optimize, test and analyze performance with varying levels of noise and a wet-lab experiment. By “stress-testing” our schema *in silico*, we were able to fit our framework to the expected noise characteristics of real Nanopore sequencing conditions, before synthesizing and sequencing *in vitro*. Our experimental validation demonstrates the practical feasibility of progressive image decoding from DNA storage, achieving low overhead while maintaining robust error correction. This work establishes a foundation for DNA-based multimedia storage systems that can provide immediate visual feedback during data retrieval, addressing key challenges in both storage density and user experience for next-generation archival applications.

